# Deep learning-based system for real-time behavior recognition and closed-loop control of behavioral mazes using depth sensing

**DOI:** 10.1101/2022.02.22.481410

**Authors:** Ana Gerós, Ricardo Cruz, Fabrice de Chaumont, Jaime S. Cardoso, Paulo Aguiar

**Author notes:** For correspondence: [ ].

## Abstract

Robust quantification of animal behavior is fundamental in experimental neuroscience research. Systems providing automated behavioral assessment are an important alternative to manual measurements avoiding problems such as human bias, low reproducibility and high cost. Integrating these tools with closed-loop control systems creates conditions to correlate environment and behavioral expressions effectively, and ultimately explain the neural foundations of behavior. We present an integrated solution for automated behavioral analysis of rodents using deep learning networks on video streams acquired from a depth-sensing camera. The use of depth sensors has notable advantages: tracking/classification performance is improved and independent of animals’ coat color, and videos can be recorded in dark conditions without affecting animals’ natural behavior. Convolutional and recurrent layers were combined in deep network architectures, and both spatial and temporal representations were successfully learned for a 4- classes behavior classification task (standstill, walking, rearing and grooming). Integration with Arduino microcontrollers creates an easy-to-use control platform providing low-latency feedback signals based on the deep learning automatic classification of animal behavior. The complete system, combining depth-sensor camera, computer, and Arduino microcontroller, allows simple mapping of input-output control signals using the animal’s current behavior and position. For example, a feeder can be controlled not by pressing a lever but by the animal behavior itself. An integrated graphical user interface completes a user-friendly and cost-effective solution for animal tracking and behavior classification. This open-software/open-hardware platform can boost the development of customized protocols for automated behavioral research, and support ever more sophisticated, reliable and reproducible behavioral neuroscience experiments.

## INTRODUCTION

Behavior is shaped by interactions between the organisms and the environment, being the most important output response of the nervous system to external (and internal) stimuli. Understanding this relationship between behavior and neural activity is the central goal of systems neuroscience, which relies on analyzing animal behavior for theorizing cognitive mechanisms and ultimately explaining the underlying neural circuits ^1–3^. Besides basic neuroscience research, the study of animal behavior plays a key role in the translational analysis of disease models, preclinical assessment of therapies’ efficacy, and also in food production industries ^3^.

The research on animal behavior has benefited from the recent technological advances in machine vision and machine learning fields, allowing for the collection and automatic quantification of vast amounts of data. Besides reducing human bias and subjectivity, and consequently allowing for the standardization of measurements across laboratories, behavioral patterns that were once unnoticed to a human observer may now be explored at different scales and resolutions ^4–6^. The first approaches to successfully combine computer vision and machine learning techniques typically relied on hand-crafted features extracted from images or video sequences that can be then used for automated behavior classification using supervised ^7–10^ or unsupervised ^11–13^ learning methods. However, such approaches are highly dependent on domain expertise for feature engineering, often losing their generalization capability in the presence of a new environment/scenario. Recent developments in the computational neuroscience field have explored deep learning techniques to meet this challenge. Most state-of-the-art systems present powerful deep learning-based solutions for pure body-part detection and tracking for pose estimation ^14–20^, but modest progress has been made for direct recognition of behavioral events ^21–23^. When compared to action detection in humans, which already achieved outstanding performance in challenging benchmarks, animals’ behavior is more complex to characterize. First, some animal behaviors are very similar to each other (more easily confused than those of humans), in which temporal information is necessary for a flawless detection (sometimes a single frame is not enough to label the behavior correctly). Recent approaches take advantage of deep architectures that integrate temporal information along with spatial information to this end ^21–23^. Also, different behaviors have different durations and temporal scales: some of them take place in long time scales, such as *grooming*, and others in short time scales, such as *rearing* or *walking*. To the best of authors’ knowledge, temporal multi-scale integration has not been explored in the context of animal behavior analysis. Another concern when planning behavioral experiments is to ensure that the environment where the animal moves is adequate to allow capturing natural behavior and yet probing for multiple parameters for its study. In particular, an important limiting factor for recording natural rodent behavior is the environment lighting conditions (which may affect animals’ biological cycle). Usually, the most natural conditions are left behind at the expense of recording conditions (higher image resolution or contrast). One possible strategy is to use cameras with infrared technology (such as deep sensing cameras). A few studies have recently begun combining deep learning methods with data from such technologies for animal behavior analysis ^24^. Finally, to effectively correlate behavioral functions with specific neural circuits, automatic behavioral analysis tools should ideally be integrated into real-time closed-loop control systems, that provide instantaneous feedback based on the current behavioral expression. There are already published tools that provide feedback control in real-time based on animal posture patterns ^9, 17, 24–27^. However, they do not satisfy all these requirements simultaneously for a complete and versatile behavioral analysis system.

Here, we introduce a novel computational solution for automated, markerless, real-time three-dimensional (3D) tracking and behavior classification of 4 classes (*standstill*, *walking*, *rearing* and *grooming*) in experiments with a single freely-behaving rodent. Combining the power of low-cost depth sensors and deep learning techniques, the proposed framework is integrated into a control platform that streams real-time mapping of input-output signals using the animal’s current behavior and position. First, we analyze the performance of advanced action recognition deep learning networks on the rodent behavior dataset. Acknowledging the importance of integrating temporal information in behavioral feature learning, we hypothesized whether abstract spatiotemporal features obtained from simple deep networks are suitable for recognizing multiple behaviors. In particular, the behavior of networks for increasing temporal extents and with multiple timescales’ branches (partially inspired in Feichtenhofer, et al. ^28^) was compared regarding their performance in detecting behavioral events. We found that temporal information from the past, using a short-time scale, is most relevant for the learning process. Second, we analyze how robust the proposed networks were at different input representations (input frame encodings, sampling rates, and resolutions), where raw depth frames at higher sampling rates and resolutions helped improve classification performance. Also, ∼21 minutes (min) of annotated video showed to be already sufficient to attain a good generalization using proposed deep networks for behavior classification. Lastly, we adapt the deep learning framework to recognize animal tracking and behavior in real-time, and we integrate it into a platform capable of closed-loop control of behavioral experiments, either for behavioral mazes or real-time drug delivery systems. Besides being non-invasive and with low latency, it provides a versatile interface to trigger different hardware actuators from either hardware sensors or behavior/tracking-dependent signals.

## RESULTS

The proposed system for online rodent behavioral recognition consists of two components: a deep learning network (**Fig. 1a**) and a real-time control module (**Fig. 1b**). The network consists of an encoder and a classifier, which is trained end-to-end. The encoder consists of two-dimensional (2D) convolutional layers, to extract local spatial features in each frame of the video sequence. The classifier is composed of a recurrent layer to learn temporal features between adjacent frames in the video sequence, and a fully-connected layers to output the behavioral classes’ probabilities (**Fig. 1a**). Networks with different architectures and input representations were studied. Whereas the deep learning network is responsible for spatiotemporal feature extraction and behavior detection, the real-time classification is used to control sensors/actuators in any maze. All these tasks can be controlled through an easy-to-use graphical user interface (GUI) for beginning-to-end management of all experiments.

**Fig 1.**
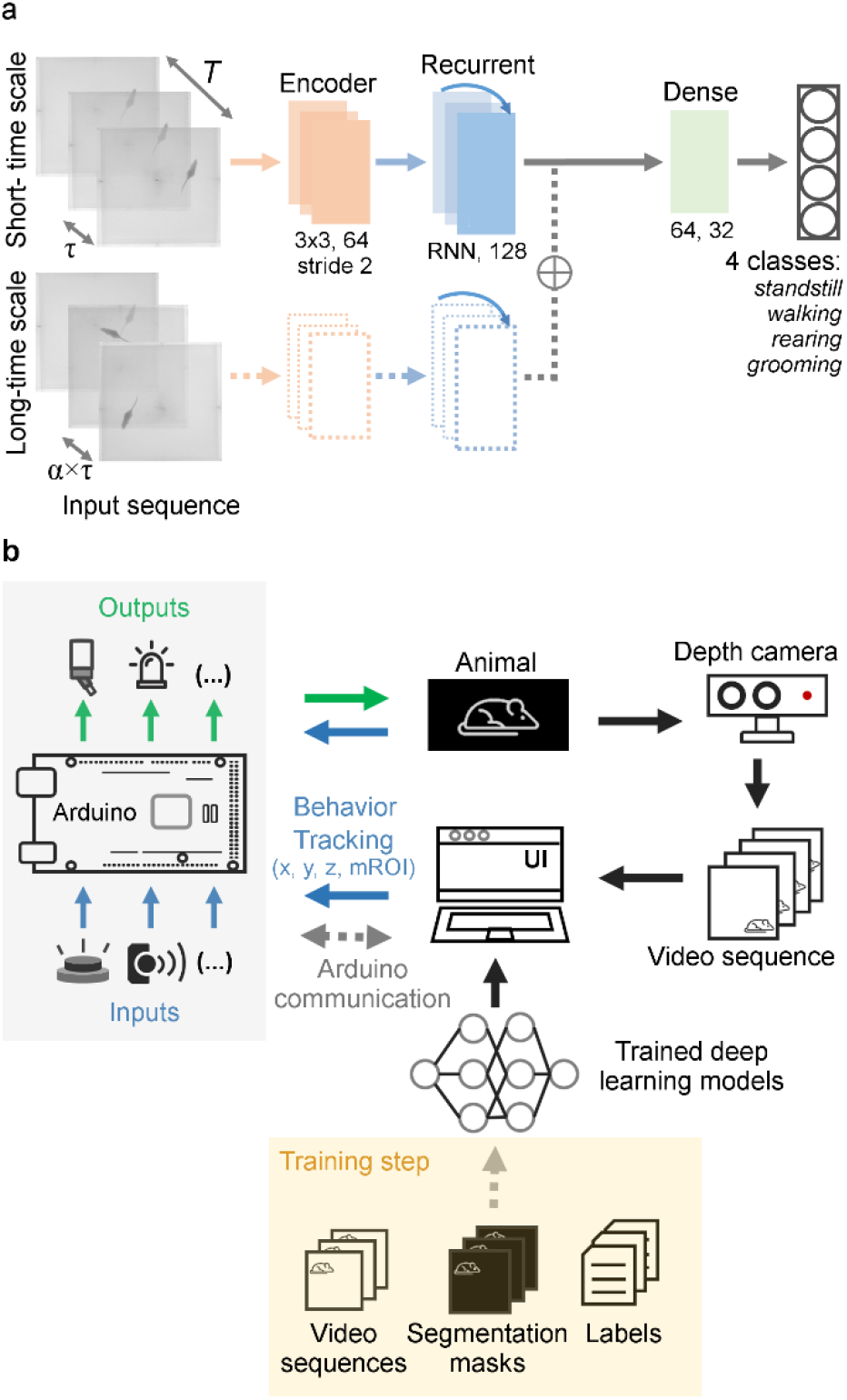
Integrated framework for the control of behavioral mazes using depth information and deep learning-based techniques. **a.** Deep learning architecture, with the two variants of the encoder, single-branch (solid line) and dual-branch (solid and dashed lines), for the automatic classification of 4 behavioral classes. Both variants receive one input sequence with a time-window of size *T* ms, with frames equally spaced over time by a temporal stride of τ. The dual-branch variant receives additionally one sequences with a different temporal stride, long-time scale pathway, that operates on a bigger time-window (α × T’) with a temporal stride of α × τ (α >1, where α is the frame rate ratio between short- and long-time scale pathways). **b.** Workflow of the closed-loop feedback system, for controlling behavioral experiments. Depth video sequences are acquired by a depth camera, and used as inputs to deep learning networks for real-time automatic classification of behavior and detection of animal’s position (x, y, and z coordinates of centroid, and any defined regions of interest inside the maze (mROI)). Such signals, together with input signals coming from any sensor hardware (blue), are sent to the Arduino microcontroller for feedback control of the actuators present in the maze (green). For real-time behavior classification and detection of animal’s position, the deep learning models must first be trained using a training set with annotated depth video sequences (segmentation masks and behavioral labels).

### Past information improves behavioral classification performance

To investigate the behavior of networks for increasing temporal extents, the time-window T of the sliding input sequences was systematically increased, with a fixed temporal stride τ=133 ms (**Fig. 2a** and **Supplementary Figure 1**). Improvements over T were observed, where models with a time-window of 10τ (approximately 1500 ms, 11 frames in the sequence) achieved the top overall results on the validation set, with a balanced accuracy of 80.0% [74.6, 83.0]%. No statistical differences were found when using as input a time-window of 4τ. The results seem to indicate that the gain of increased time-window is clearer for networks with a smaller time-windows, with a converging trend towards time-windows above 1000 ms. This is aligned with the timescale for the analyzed animal behavior classes (where the timescale for variation is in the order of 1 second) (**Fig. 2b**). For time-windows smaller than 300 ms, the performance significantly dropped. When no temporal information was taken into consideration, using a model with only one input frame, the lowest overall accuracy was achieved, as well as category F1-score, showing that not only spatial information within a particular frame may be important but also its motion content across different frames. In fact, when performing manual annotations, ethologists often need to double-check previous frames to annotate the current one, which also seems to happen in these networks.

**Fig. 2.**
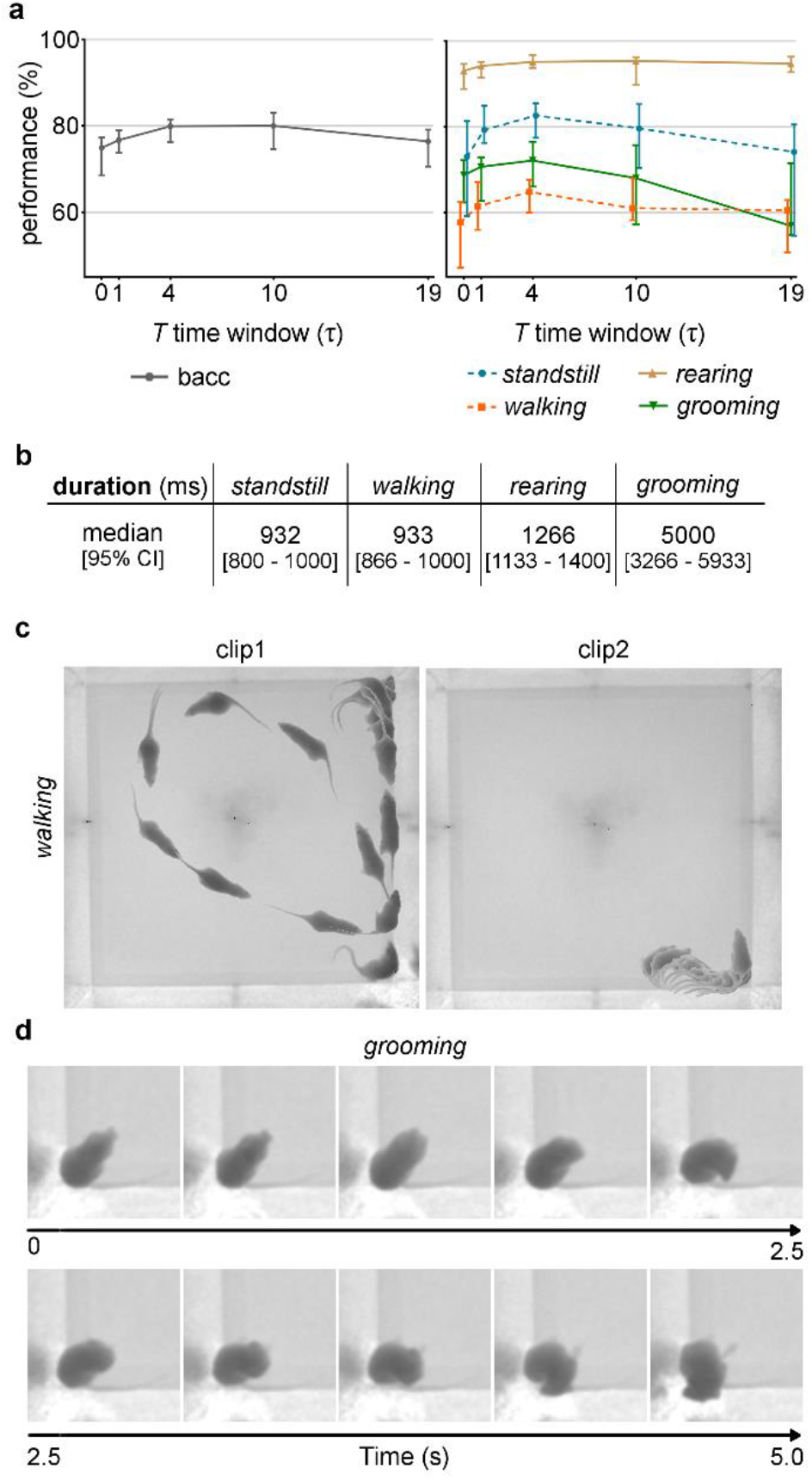
How much temporal information does the network need for rodents’ behavioral learning? a. Results using single-branch architecture of varying temporal extents. Left: Overall balanced accuracy (bacc) for increasing temporal extents. Right: F1-score per class. Time window *T* in units of τ (τ = 133 ms). Data represented as median ± 95% confidence interval (N = 5 trials). **b.** Behavioral events’ duration, in milliseconds (ms). Data represented as median ± 95% confidence interval. **c.** Stroboscopic montage in which each animal position represents raw depth frames extracted at every 266 ms for 2 different *walking* clips. **d.** Sample clips with frames extracted at every ∼500 ms, for a single *grooming* clip.

Out of all 4 classes, no behavioral event has a monotonic decrease with the increasing temporal extent, and overall their recognition seems to benefit from time-windows smaller than 1000 ms (category F1-score systematically increasing over ***T***, until approximately 1000 ms). This effect is particularly clear during *standstill*, *walking* and *grooming* events, where F1-score performance seems to slightly decrease for time-windows greater than 1000 ms. In fact, *standstill* and *walking* are events that usually last for a shorter period of time, compared to other behavioral events, containing approximately 932 [800 – 1000] ms and 933 [866 - 1000] ms as median duration (**Fig. 2b)**. For this reason, they do not seem to benefit from long time-windows for accurate recognition. Furthermore, *walking* is the class with the lowest overall performance and one possible explanation could be the fact that *walking* is the class containing greater intra-class movement variability (either in terms of complexity of geometric shapes, sequences’ durations and movement speeds) (**Fig. 2c**). The behavioral event that appears to be the most sensitive one to increasing the temporal extents is *grooming*. Using manual annotations given by the ethologists, this action is typically composed of several stationary periods interspersed with shorter periods of movement, in which the animal changes its position momentarily without leaving the *grooming* event. Long-term networks, with larger time-windows, can, thus, easily confuse *grooming* with *standstill* events (not shown), due to this heterogeneity within one single *grooming* sequence (one example is shown in **Fig. 2c**, where a sequence of *grooming* frames was sampled at every 500 ms). On the other hand, *rearing* is the class with the highest performance for the different time-windows studied, not seeming to benefit from the increase in temporal extents. In fact, this is the less ambiguous behavior in the current classification task, because of its easy-to-distinguish geometric shape and lower depth values, and usually it is enough to analyze closer frames to confirm it.

### Short-time scales are the most relevant for the learning process

Additionally, two variants of network encoder, single- and dual-branch, were systematically compared to study the impact of having temporal information of different scales. While in the standard single-branch networks the input is a time-sliding sequence of size *T* ms, with a fixed temporal stride τ ms between frames, this dual-branch network is fed with input sequences with different temporal strides in each pathway, as a way to understand if having multiple time scales helps in the learning process (**Fig. 1a**). The idea is for the two pathways to exploit temporal information of a different scale: the short-time scale provides information hidden in temporally neighboring frames, giving clues about animal’s movement at fast temporal changes, while the long-time scale may help distinguish between different behaviors at slower temporal changes (namely, transitions between behavioral states).

To allow direct comparison, a single-branch architecture, with a time-window of 2τ and a temporal stride of 133 ms, and a dual-branch architecture, with different frame rate ratios *α* between the short- and long-time scale pathways, were trained and validated. The single-branch and dual-branch *α* = 5 appear to have similar overall performances (**Fig. 3a**), even for per-class recognition; however *α* equal to 10 (which means doubling the time-window for that pathway) seems to decrease performance. These results are in line with the conclusions of the previous section, where behavior learning does not seem to benefit from very distant temporal information (irrelevant frames are being taken into consideration, degrading network’s performance).

**Fig. 3.**
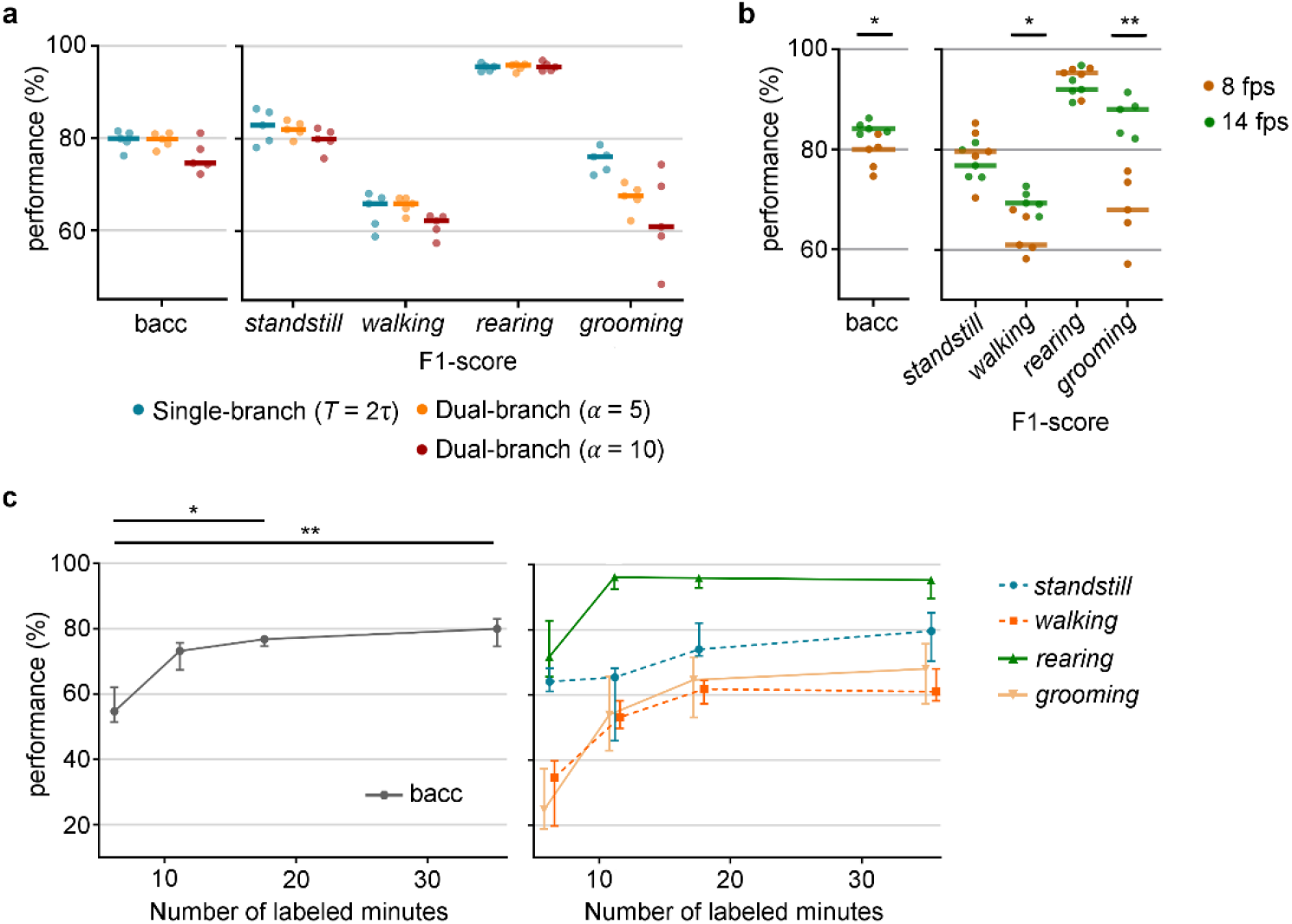
a. Which time scales are most relevant for the learning process? Comparison between architecture with different temporal scales: single-branch and dual-branch (*α* = 5 and *α* = 10), regarding overall balanced accuracy (bacc), and F1- score per class. **b.** How should time be distributed to increase performance? Comparison between different temporal strides τ between adjacent frames (τ ∈ {67, 133} *ms*, corresponding to approximately 15 or 8 frames sampled per second, respectively). **c.** How much information does the network need to learn? Overall and per-class classification performance as function of number of labeled minutes. Data represented as median ± 95% confidence interval (N = 5 trials). * *p* < 0.05; ** *p* < 0.01. Statistical analysis only for overall balanced accuracy for the sake of readability. Additional statistical analysis on **Supplementary** Figure 2.

### Different input sequence’s representations improve networks’ learning

To further understand whether the temporal extent of video input sequences or their sampling frame rate with which the network is fed has more impact on learning rodents’ behavior, networks with different temporal strides τ, but a fixed time window *T =* 10τ, were also compared (**Fig. 3b**). Significant improvements were observed when using higher frame rates (smaller temporal strides), with an increase of approximately 5% in the overall performance (with a frame rate equal to 15 fps, the median balanced accuracy reached 84.1% [83.0 - 86.2]%). In particular, *walking* and *grooming* events greatly benefit from increasing the input frame rate. This could indicate that a higher temporal resolution is needed to detect movement oscillations inherent to these types of heterogeneous behavioral events.

As part of the networks’ systematic study, the effects of input resolution and input depth encoding were also examined. The highest resolution (256x256) achieved the best results, with an overall performance of 85.9% [82.8 – 86.6]%. All behavioral events seem to benefit from increased resolution, in particular *grooming*, with an increase of approximately 44% over the lowest resolution (**Supplementary Figure 3A**). When changing input depth encoding, networks trained with raw depth frames outperformed any other depth encoding techniques, with surface normal inputs reporting the worst performance, yielding an overall accuracy of 71.8% [60.9 - 75.8]% (**Supplementary Figure 3B,C**).

### High performances achieved with a reduced training dataset

In order to determine the approximate amount of annotated training data required for good network performance, the size of the training set was systematically varied (**Fig. 3c**). As expected, overall performance increases for increasing number of training images. Even 10k labeled frames (approximately 21 min of labeled data) were enough to achieve a good generalization, above 70%, with performance degradation in *walking* and *grooming* events. In fact, the effect of changing training size is most significant in these classes, where increasing 20 min of annotated data leads to a gain of almost 45% in per-class performance. Peak performance was reached with 30k training examples (corresponding to approximately 1hour of labeled data).

### Behavior is accurately detected in unseen depth videos

The behavior of the network against a completely unseen testing set is the ultimate study to quantify recognition performance and generalization capability of the model (**Fig. 4 a,b)**. After being trained with the best set of parameters, the model achieved an overall accuracy of 82.2 % [78.5 – 83.9]%. Together with the ethograms automatically generated (**Fig. 4b**), these results indicate that the proposed automated classification method captured the overall patterns of behavior in the new videos.

**Fig. 4.**
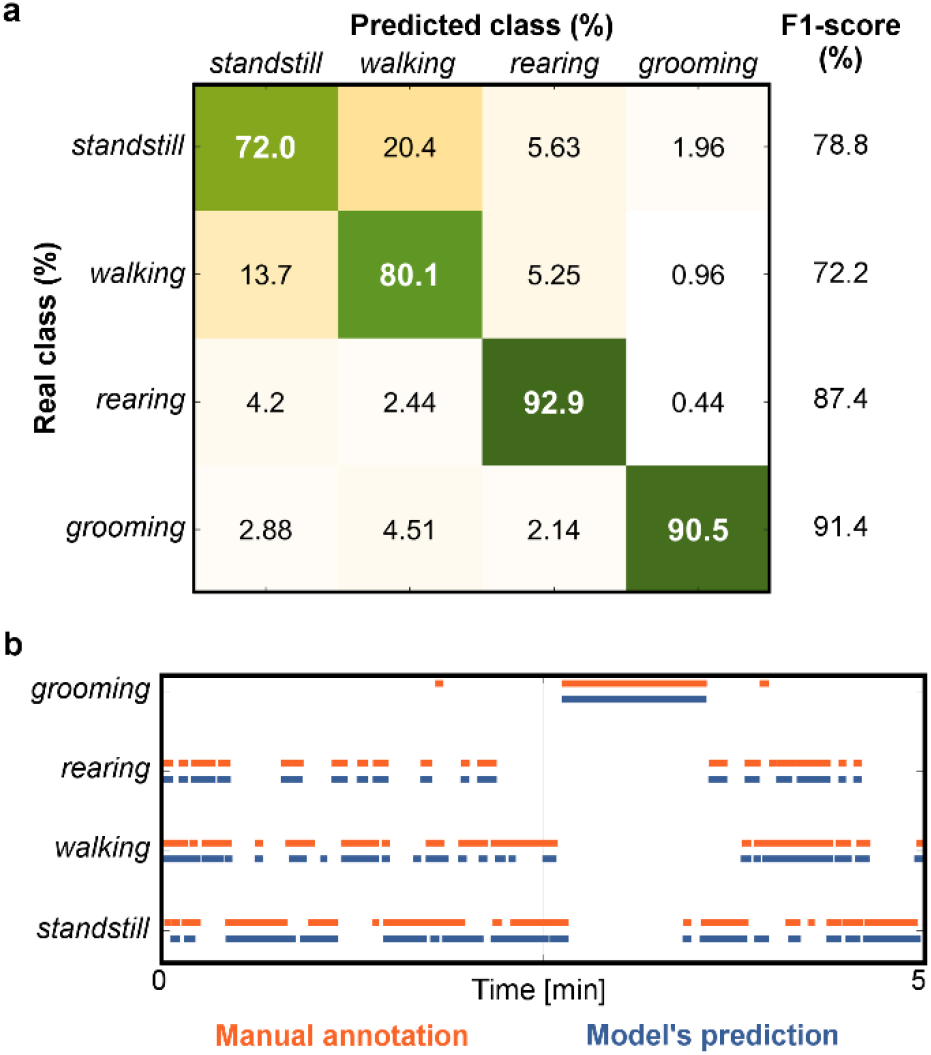
How does the best network behave for an unseen test set? a. Example of normalized confusion matrix for a detailed analysis of automated behavior recognition errors, and corresponding F1-scores for each class. **b.** Example of ethogram for a comparison between automated model’s detection (orange) and manual annotation (blue), over 5 min of testing video.

Regarding per-class performance, *rearing* is the behavioral event with the highest performance, attaining 87.2% [86.0 – 91.1]% F1-score, in accordance with previous results. Also, *walking* periods belong to the most misclassified behaviors, which are occasionally classified as *standstill* events (example in **Fig. 4a**), given frames’ heterogeneity on shape and speed.

### Closed-loop system achieves low-latency feedback based on animal behavioral/tracking patterns

In order to create a system capable of controlling a behavioral task based on animal behavior/position, it is necessary to close the loop between automatic detection of behavioral events and experimental operant conditioning hardware. A control platform, combining a depth-sensor camera, computer and Arduino microcontroller was constructed to allow mapping of input-output control signals using the current deep learning detection of animal behavior and position. Additional results on the performance of the segmentation task using deep networks can be found in **Supplementary Figure 4**. To demonstrate the applicability of the closed-loop framework in triggering signals based on animal behavior, an experiment was designed in which four actuators (in this case, LEDs) were turned on when the rat performed one of the four behavioral events: *standstill*, *walking*, *rearing* and *grooming*. The behaviors and tracking positions were automatically detected by previously trained deep networks, that, together with input signals coming from different sensors, are sent to the Arduino board to control the output devices. This setup achieved delays from image acquisition to detecting the behavior+tracking position (image-event delay) as fast as 28.9 ms [26.95 – 31.86] ms, for an input resolution of 128x128 (**Fig. 5a**). For larger images (256x256), the delay increased about 8.9% (full results from additional configurations can be found in **Fig. 5a**). The proposed system, with the advanced hardware configuration (GPU settings) and for the smaller resolution, reached a performance time of 32.9 ms [32.8 – 34.9] ms from predicting one behavioral event+tracking position to the next one (event-event delay), including Arduino output generation, frame acquisition and processing, and behavior/tracking position detection. Finally, sending the signal to the Arduino board and sending back the signal to the computer took an additional 0.457 ms [0.457 – 0.460] ms, when compared to just turning on the LED – event-LED delay (0.914 ms [0.913 – 0.914] ms). Thus, the Arduino response is not constraining the runtime from event detection in one frame to the next frame, and it can be almost entirely attributed to intrinsic camera frame rate, behavior/tracking detection and additional processing.

**Fig. 5.**
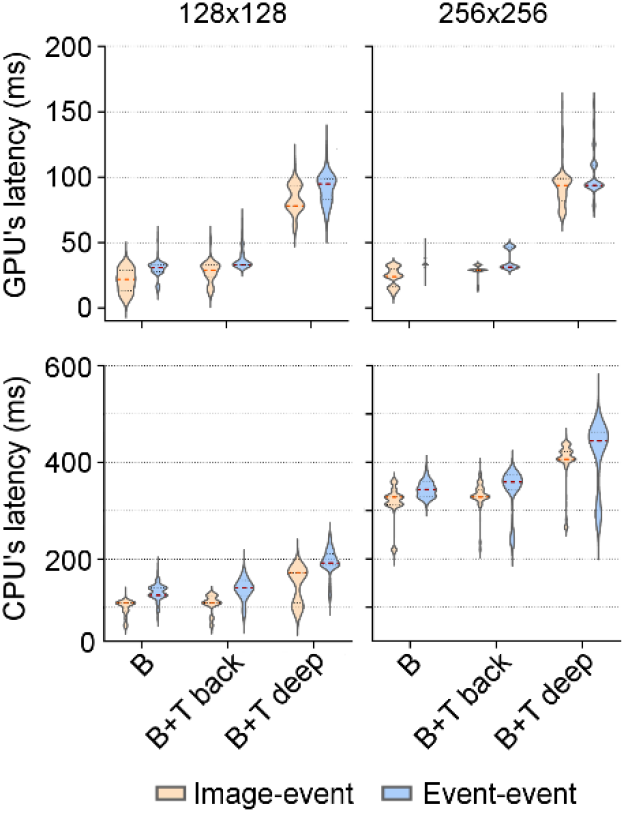
How to close the loop for behavioral experiments? Latencies, in milliseconds (ms), from image acquisition to obtaining an event (image-event) and from the last event detected to the current event detected (event-event), using CPU or GPU processing. Latencies were estimated for automated predictions of behavior only (B), behavior and tracking using the background subtraction method (B + T back), and behavior and tracking using a deep model-based method (B + T deep). The width of the violin plots represents the probability density of the data, with the median and 95% confident interval represented as red and black dashed lines.

### User-interface allows end-to-end control of behavioral experiments

Acknowledging the importance of embedding all algorithms in a user-friendly application suited for reasearch environments, we developed a full-featured, easy-to-use and freely available software interface (**Fig. 6a**), requiring no programming by the end-user.

**Fig. 6.**
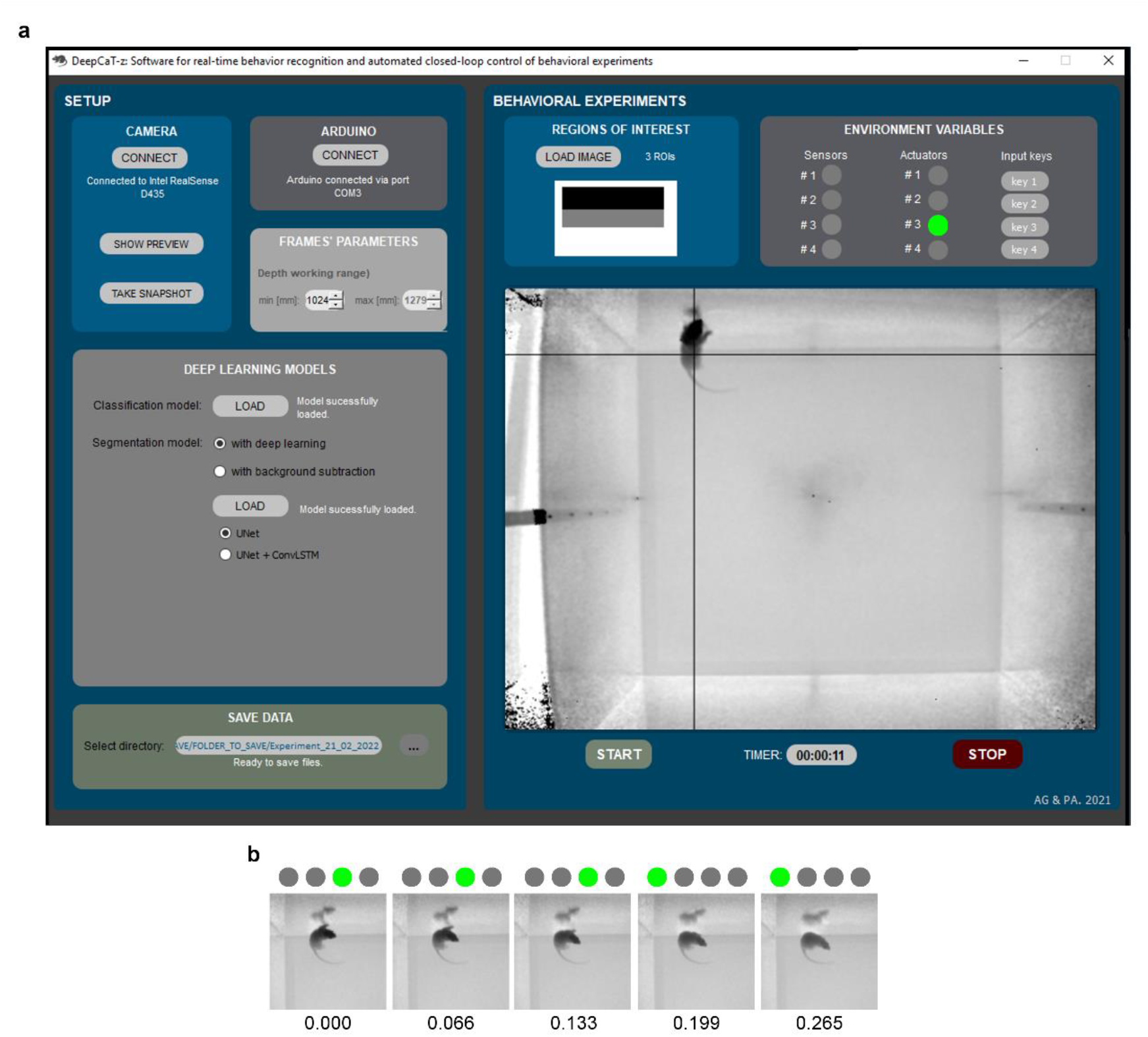
How to easily control behavioral experiments? a. Graphical user interface for automating real-time closed-loop behavioral experiments. **b.** Example of a *rearing* followed by a *walking* sequence, with corresponding LED status (as it appears in the graphical user interface), from the test video sequence. Image timestamps in seconds are presented at the bottom of each image.

Behavior classification and/or tracking are performed using different available methods, chosen by the user, and detected using uploaded trained models. The GUI provides online information regarding hardware modules states, animal’s behavior and position, allowing full control of the entire system. In particular, the state of 4 sensors and 4 actuators are updated in real-time, in which a LED-type icon is turned on upon the first image in which a behavioral pattern was detected, and subsequently turned off upon the first image in which the pattern is no longer detected (**Fig. 6b**). This allows for a fully closed-loop stimulus’ framework. The GUI also includes an option for users to upload an image containing ROIs for a more versatile and complete behavioral analysis. All useful information recorded during the experiment (depth frames, tracking and behavioral classes’ information with sensors/actuators states for each timestamp) can be exported to a user-defined directory for further analysis.

Overall, a cost-effective and easy-to-setup framework was created. The entire system consists of a computer running the GUI, connected to a depth camera (e.g., Intel RealSense Depth Cameras, of ∼300 €) and an Arduino (e.g. Mega 250, of ∼35 €). Sensors and actuators can be directly connected to the Arduino board, and the quantity and type depend on each experiment’s goal. The source code of the software, together with the user-guide manual, list of hardware materials and video examples, are publicly available for download at GitHub (https://github.com/CaT-zTools/Deep-CaT-z-Software).

## DISCUSSION

We have presented a fully integrated framework that can provide real-time feedback based on automated rodents’ behavior classification and tracking position, using specialized deep neural networks to extract information from frames acquired with depth-sensing technologies.

With the developed algorithms, we demonstrate that cutting-edge deep learning models can be used to learn features from depth video sequences, without the need for feature-engineering approaches. In fact, this is one of the main reasons why deep learning-based methods can be more powerful than conventional behavior classification ones, avoiding user bias in the learning process and allowing for more easily tunable and generalizable systems. This is particularly important in basic research where environmental setups or animals’ appearance/strains may be changed depending on the objectives of each experiment and yet it is possible to successfully apply the same methods ^3, 6^.

Furthermore, the capabilities of these deep learning networks were extended to learn feature representations exclusively from depth information. Although several deep learning-based studies have been published using depth frames for detecting human behavior, depth information is usually incorporated using multi-branch architectures, combining color and depth inputs from multiple streams for motion capture ^29–31^. Here, we focused on depth images and how information can be successfully retrieved for animal behavior extraction. Analyzing behavior with only depth information has four important advantages. Since these frames are acquired by infrared sensors, videos can be recorded in dark conditions (where color information is useless) without disrupting animals’ natural behavior (mainly in nocturnal animals, such as rodents). Also, with this technology, color contrast between the animal and the background is no longer a problem for detection/tracking purposes. Conventional methods usually use markers or methods dependent on animals’ color coating ^32–36^, which can be avoided using depth-sensing information. In addition, 3D information can be retrieved from a single camera, and so setting complicated stereo-vision setups is no longer needed. Finally, to further facilitate the int*e*gration of computational methods in the laboratory and industry fields, low-cost acquisition devices are required, combined with good performance and, at the same time, quick data acquisition and low computational cost.

Therefore, the use of depth technology, such as *Kinect*-based cameras, showed to be an alternative strategy to be applied in behavioral experiments. Since there are no state-of-the-art studies exploring the use of depth information in the context of feature extraction for animal behavior classification, we also perform a systematic study to understand the best ways to represent network inputs and how we can improve models’ performance. By using deep learning networks that incorporate spatiotemporal features, it was possible to conclude that temporal information is very relevant for learning animal behavioral patterns, especially in some classes (*standstill* and *walking*, which contain a strong dynamic component). These results are in agreement with the fact that temporal information of video data can provide additional clues hidden in temporally neighboring frames for the recognition of actions/behaviors or segmentation of frames ^29, 37^. By using a fixed temporal stride between input frames of approximately 133 ms, the performance of networks is significantly improved for input video sequences with a time-window of approximately 1.5 seconds. As expected, some animal behaviors are of very short duration, with rapid transitions, sometimes imperceptible by humans, and for this reason, deep neural networks for animal behavior classification must be carefully designed to support finer temporal analyses. In addition, results showed that neither long-time scales nor multi-scales seemed to be advantageous for detecting animal behavior. One possible explanation is that long-time scales include frames too far apart in time, containing irrelevant information to learn useful feature representations for the current frame. Although with our system we didn’t see advantages in the multi-scale analysis, we hope that it can be further explored in the context of animal behavior. For example, in a system with higher frame rates, it may be useful to also explore shorter time scales.

Along with the fact that higher resolutions and higher sampling rates in raw frames (without preprocessing or encoding) significantly improve the performance of proposed deep networks, the results give an insight on how to build, train and fine-tune networks to better learn rodent behavior using depth-sensing information. Finding that ∼21 min of annotated videos are already sufficient to achieve high generalization rates strengthens the contributions of the proposed system since a core goal of automating the analysis of behavior is reducing the manual annotation effort. In this sense, once the deep learning model is trained, the system is ready to assist in any behavioral experiment without additional user-time, allowing for more reproducible results and reducing variability imposed by inter-human annotations. Recent works have made some progress toward the goal of supervised classification of rodents’ behavior using deep learning techniques to improve conventional feature-engineering-dependent methods. Marks, et al. ^22^ developed *SIPEC:BehaveNet* for behavior recognition, which was tested in a dataset acquired with a conventional camera and containing freely behaving mice whose behavior was labeled with only 3 classes ^38^. Although claiming superior performance to Sturman, et al. ^38^ proposal, *SIPEC:BehavNet* achieved lower overall performances for *supported rearing* and *grooming* events (mean ± standard error of the mean: 0.84 ± 0.04 and 0.49 ± 0.21, respectively), when compared to what we were able to report here. *DeepEthogram* is another recent tool for frame-based classification of animal behavior in RGB videos ^39^. High overall performances (overall accuracy) were obtained for datasets containing mice behavior with more than 4 classes. However, performance per-class (F1- score) is substantially impaired for some behaviors, in particular, the rarest and most challenging behaviors in the dataset (average F1-score above 70%). This shows evidence that attention must be paid to metrics performance when dealing with highly unbalanced datasets. Overall, both methods fall behind some strengths that our method shows, needing more than 70 min of labeled data to achieve a comparable performance (overall accuracy above 70%) and not being suitable for natural environmental conditions in the analysis of rodents’ behavior.

In order to improve the potential of the proposed system and create an integrated tool that would boost future development in understanding behavioral patterns and neuronal activity relationship, deep learning-based detection of behavior was used to provide event-triggered feedback in real-time. The loop between animals’ maze, depth frames acquisition, and automatic streaming of behavioral patterns was closed using input and output devices connected to an Arduino microcontroller. From detecting one behavioral event to the next event in a consecutive frame, the system was able to achieve real-time feedback control, with latencies of less than 33 ms with GPU-based configuration. These results are below the frame rate of the camera used (which typically is reduced to ∼15 fps in low light conditions), and so, in theory, more powerful infrared cameras could be tested. Research on developing real-time applications for neuroscience research has been advancing in recent years. However, efforts have essentially been directed towards tools to detect animal’s posture, rather than classifying directly the behavior. Both Forys et al., 2020 and Schweihoff et al, 2021 developed software and hardware to enable real-time estimation of mice posture, and achieved latencies of 30ms using comparable computational configurations, from frame acquisition to detecting a posture of interest (slower image-event delay than what we were able to achieve) ^17, 26^. Kane, et al. ^25^ reported higher computational performances for the same task, with a 16ms delay from image-LED event (for equivalent image resolution and hardware configurations). However, it is worth emphasizing that our 30-fps figure is achieved when both behavior classification and tracking position are available, which gives the tool versatility for different research applications. To the best of authors’ knowledge, Nourizonoz, et al. ^24^ were the first to try to detect animal postures as well as simple behaviors in naturalistic environments, using multiple cameras with infrared-based technology. Real-time detections were achieved to enable reinforcing a simple behavior (*rearing*) by operant conditioning. Although with high performance in naturalistic environments and taking the first steps in moving forward to correlate posture with neural circuits by optogenetics stimulation, the detection of a single behavior from posture was achieved using a set of geometrical rules. This approach may not be sufficient to classify more sophisticated behaviors, or computationally heavier when classifying multiple behaviors.

A key aspect of the design of the whole system is its versatility and how different modules can be adapted to different research goals. In particular, several tracking algorithms were made available, depending on model’s performance and computational power. This flexibility may be important when real-time detection is not required but offline high-performance detection is needed. Also, many sensors and actuators can be easily adapted to the Arduino microcontroller to finer control of animal’s maze, and the automation control code is prepared to be further extended. Even so, recent advances in multiple animal behavior analysis and tracking ^9, 16, 40^ could be included to further enhance this versatility. System adaptation is, in theory, straightforward, however, the triggers for feedback control need to be carefully designed when dealing with complex social behavior. Furthermore, the list of behavioral events/classes can be further extended. Here, the potential of deep neural networks can be explored, since they are able to extract relevant features without the need for feature re-engineering, unlike conventional machine learning methods.

Taking all the contributions together, we believe that the flexibility and yet easy-to-use characteristics of this real-time feedback framework may open the door to further studies and broader applications, allowing more high-throughput and rigorous behavioral experiments while less invasive for laboratory animals.

## MATERIALS & METHODS

### Dataset

An open-access RGB-D behavioral dataset, available at https://doi.org/10.5281/zenodo.363613510, was used for all experiments. Details on the experimental procedures, video acquisition and manual annotation of rodent’s behavior can be found in ^10^. In brief, the dataset is composed of 10 to 15 min RGB-D video sequences of individual Wistar rat behavior, recorded with a *Microsoft Kinect v2* camera (512x424 depth pixel resolution). The maximum frame rate is 30 frames per second (fps), but this value typically drops to 10 to 15 fps in low light conditions. A subset list of classes was considered here with the four most commonly used state behavior states: *standstill*, *walking*, *rearing* and *grooming*. A randomly selected subset of these fully annotated recordings was considered for the experiments and denoted as *dataset-100k* (∼2.20 h in 26 subvideos, approximately 100,000 frames total, with a time difference between two consecutive frames of approximately 67 milliseconds (ms)). Only the depth frames were kept for analysis.

### Proposed deep learning model Architecture

Two variants of the encoder were considered – the single-branch and the dual-branch. In both architectures, frames are individually encoded by four 2D convolutional layers (64 filters, 3x3 kernel size, 2x2 stride, rectified linear unit (ReLU) activation). After the encoding part, a recurrent layer (RNN, 128 hidden state features) takes as input the sequence of spatial features output by the feature extractor and integrates it over time for both temporal and spatial dynamics learning. Two fully-connected layers (64 and 32 filters) and a softmax output layer are used for the final recognition of behavioral classes. In the case of the dual-branch, both pathways work on different time-windows: the short-time scale pathway receives as input a pre-defined time-window *T’* with the same temporal stride τ as the single-branch network; the long-time scale pathway operates on a bigger time-window (*α* × *T*′) with a temporal stride of *α* × τ, where *α* > 1 is the frame rate ratio between short- and long-time scale pathways. Two recurrent layers are used for each branch, which are then concatenated before the fully-connected layers.

Since recognizing rodent’s behavior is a challenging task, either due to the size of the animals or the nature of the behaviors (faster movement, higher similarity and greatly dependent on temporal information to be clearly distinguished), the feature extraction process needs to be carefully designed to avoid confusion between behavioral events. For this reason, 2D convolutions were chosen, instead of the currently used 3D convolutions for spatiotemporal learning, in order to process spatial and temporal content separately and thus avoid mixing information of different scales. The reduced number of convolutional layers and the number of filters at each layer allow the entire network to be computationally lightweight and capable of being used for real-time inference afterwards.

### Training

The models were trained from scratch using the Adam optimizer, with a batch size of 16 video sequences with a time-window of *T* ms, and a learning rate of 1 × 10^−4^, for 100 epochs. A dropout layer was used before the recurrent layer, with a dropout ratio of 0.5.

Initially, the dataset was split into training (70%), validation (10%) and testing (20%) sets that are maintained throughout the experiments. The validation set was used to compare the performance of different models when performing ablation studies. To address the problem of having a highly imbalanced dataset (*standstill* 40.3%, *walking* 28.7%, *rearing* 11.7%, and *grooming* 19.3%), the video sequences of each class were oversampled until their frequencies were uniform.

## Experiments

For a systematic study of networks’ performance, the effect of increased temporal information was evaluated, by changing different parameters in each experiment. First, the impact of changing the time-window *T* of the input sequence was tested, with *T* ∈ {0τ, 1τ, 4τ, 10τ, 19τ} *ms*, corresponding to a network input with 1 (single-frame), 2, 5, 11 and 20 frames in total, respectively, sampled with a fixed temporal stride τ of 133 ms. Also, the temporal stride τ between adjacent frames (τ ∈ {67, 133} *ms*), was also evaluated, which corresponds to approximately 15 or 8 frames sampled per second, with a fixed time-window. Finally, the frame rate ratio *α* between short- and long-time scale pathways for the multi-branch architecture (*α* ∈ {5, 10}) was varied. These temporal parameters were chosen in order to make the network responsive to the different behavior timescales present in the original dataset. In this sense, and taking into consideration the camera’s frame rate, the capability of the network of capturing both fast behavioral events (in the order of a few hundred milliseconds) and slower events (in the order of a few seconds) was explored. Also, different spatial resolutions of {64, 128, 256} pixels and input encoding modalities were tested. Besides raw 8-bit depth frames, depth jet-encoding ^41^ was applied to depth frames, in which the depth information is distributed according to the jet colormap, transforming the one-channel depth map to a three-channel color image. Also, surface normals were used to encode the depth frames into a three-channel image representing form and surface structure (implementation details in ^42^). Unless otherwise noted, the full *dataset-100k* was considered for analysis, and the default parameters for the systematic study were: *T* = 10τ, τ = 133 ms, spatial resolution of 128 pixels in raw depth frames. The influence of training set size on network generalization was also benchmarked. Different training sizes were selected and each subsampled training set was used to train the network, and compared with the same validation set (using the default parameters’ set as well).

### Data augmentation

To improve the robustness and generalization of the models, data augmentation was performed with random perturbations of the training set during training, that included: full-rotation around the center (90/180/270°); horizontal flipping; resized cropping and brightness variation (by sampling an additive value from a uniform distribution, [-0.15, 0.15]). As the input of all models is a frame sequence of approximately *T*⁄τ frames, the same augmentation operations were performed on each frame in this set.

### Model evaluation and metrics

The validation set was used for models’ comparison and evaluation, and all analyses reported share the same validation set, for a total of 5 runs for each experiment. The hold-out testing set was further applied to evaluate the performance of the best-chosen model to an unseen set. To evaluate the overall performance of the different proposed methods, balanced accuracy (average of recall obtained on each class) was calculated. Performance per class was assessed using confusion matrices and corresponding F1-score.

The F1-score is the harmonic average of the precision and recall, calculated as follows:

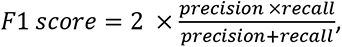

where 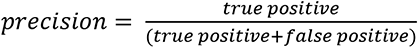 and 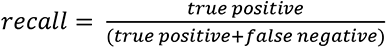. These metrics are better suited to deal with imbalanced datasets.

### Real-time control system

The entire control system consists of software and hardware modules configured to create an automated closed-loop tool. It is made of five main components: the control computer, the interface board, the control software, the video camera and the maze hardware modules (**Fig. 1**). Frames acquired by a depth camera are fed into the trained deep learning models, which will automatically detect both behavioral events and the animal’s position in the maze. The network outputs are sent to the interface board that, together with existing sensor outputs (e.g., buttons, maze sensors), controls circuit actuators (e.g., maze feeders, light-emitting diodes (LED)s). The computer is used to operate the entire circuit by a graphical user interface (GUI), either sending messages to the interface board or acting directly on the maze hardware modules.

### Interface board

An Arduino microcontroller (Mega 2560) was used as the interface board between the computer and the hardware modules, and the communication is established using a communication (COM) port. The microcontroller board has 16 MHz clock speed, and 54 digital input/output ins that can be connected to different maze hardware components, such as animal feeders, LEDs, maze sensors, and buttons. After being connected to the computer, the Arduino board communicates via Arduino integrated development environment (IDE). The user writes the Arduino code for the automated control in the IDE, uploads it to the microcontroller which executes the code to interact with the input and output hardware modules. Notice that, once uploaded, the code can run regardless of the connection between the Arduino and the computer.

### Control software

The automated control software consists of the following components: the automation control code, the trained deep learning models for detection, and the data acquisition and communication protocol.

#### Automation control code

Arduino code is written within the Arduino IDE (in a language very similar to C++) and it was carefully organized to segregate the code for specific logic state implementations (automated control) from all other maintenance code (such as reading and writing data to the communication port (COM). To do so, a specific user-defined function was created, which has access to all critical variables for the control, such as sensors’ and actuators’ states, and animal’s position and behavior. Inside this function, the user can easily define the conditions of stimuli-response that characterize each behavioral test experiment.

#### Deep learning models

In order to automatically classify the behavior and calculate the position of the animal using deep learning methods, previously trained models are imported and directly used for predictions. For the automatic classification of behavior, the single-branch model was trained according to the protocol previously described (input sequence of raw depth frames, with a time-window of approximately 1330 ms, acquired at a frame rate of 15 fps). For the estimation of animal’s position, two different methods were made available to the user: deep learning-based model for semantic image segmentation, and conventional background subtraction model, both followed by centroid calculation. The deep learning-based model combines two ingredients from deep networks’ knowledge in order to perform semantic segmentation taking into consideration temporal information: U-Net model as backbone architecture, and (optional) convolutional Long Short-Term Memory (ConvLSTM) layers, learn spatiotemporal features. The traditional U-Net architecture was reduced to only one convolutional layer per block, fewer filters per layer (32) and it was extended by placing two ConvLSTM layers, one between the encoder and the decoder, and the other one before the last dense layer (different positions in the network, as well as different architecture parameters, were tested to ensure maximum performance yet reduced inference time and memory (**Supplementary Figure 4**)). The network was trained from scratch using 1220 train and 320 validation video sequences (previously annotated to obtain the segmentation masks), with ADAM optimizer and dice binary cross-entropy loss function.

A conventional background subtraction method was integrated in parallel to provide a computationally lighter alternative yet with lower performance (mainly in frames with dynamic backgrounds). Using this method, the segmentation mask containing animal’s pixels is produced by subtracting the present frame with the background model (frame of the behavioral experimental setup without the animal). From the segmentation mask, the position of the animal is calculated as the centroid of the detected object/animal. For details on algorithm’s design and performance, please check Gerós, et al. ^10^.

For a more complete information about animal’s movements inside the maze, the system allows the user to define spatial regions of interest inside the maze (mROI), by uploading an image file with the same resolution as the acquired frames, with the different mROIs painted uniformly with different colors. Those regions are automatically detected after getting animal’s tracking, and they will be used as input for the Arduino board to control the hardware mazes, if needed.

#### Data acquisition and communication

To establish the communication between the COM port and the Arduino board, a communication protocol was defined. The computer communicates with the interface board by sending the behavioral classification, tracking and mROI outputs (as well as a flag for any keypress), in the form of a characters’ list separated by commas. Each character encodes information for the behavioral state (S for *standstill*; W for walking; R for *rearing*, and G for *grooming*), tracking (x, y and z coordinates of the centroid), mROI and a key-pressed flag (both encoded as integers). On the other hand, the Arduino board sends information regarding the status of each of the sensors and actuators (binary coded, on/off) back to the computer.

### Video camera

The acquisition protocol was developed using a new generation of low-cost depth cameras, the Intel® RealSense Depth Cameras (in particular, D435 model), acquired with 512x424 depth pixel resolution and at a maximum of 30 fps.

### Computational performance: inference and latency times

To test time-performance of the system, a video of a freely-walking rat was used to simulate a camera feed from an animal in real-time, and single frames from the video were loaded at the maximum rate of 30Hz. The bidirectional communication with the Arduino board was achieved from either four input sensors and signals from the computer, and four output actuators (in this case, LEDs). Three latency periods were measured: (a) the delay from image acquisition to detecting the behavioral state/tracking position (image-event delay); (b) the delay from detecting one behavioral event/tracking position to the next event/tracking position (event-event delay, including *Arduino* response, mROI detection, GUI updates and saving images to external folder); (c) the delay between sending a behavioral state to the Arduino and turn on the corresponding LED (event-LED delay, with and without output feedback of Arduino). The first two latency times were determined using software timestamps and the last one was measured using the oscilloscope.

### Computing hardware

All experiments, including inference speed and feedback control tests, were conducted on an Intel Core i9-7940X (128 GB RAM), and a NVIDIA GeForce RTX 2080 graphics processing unit (GPU) (8 GB RAM), running *Windows* 10, with *Python* 3.9 using *PyTorch* (1.8.1) and *TensorFlow-GPU* (2.5.0) frameworks. All algorithms were integrated into a user-friendly GUI, designed in the *Qt Creator* (*The Qt Company*, Finland) environment and implemented in *Python* language.

### Statistical methods

Statistical analysis was performed using GraphPad Prism version 7.00 (GraphPad Software Inc., CA, USA). The method of D’Agostino & Pearson was used as a normality test, and parametric or non-parametric tests were chosen as appropriate. Statistical significance was considered for p < 0.05. Parametric data are expressed as mean ± standard deviation (SD), and non-parametric data are expressed as median and 95% confidence intervals.

## Data availability

## Code availability

The source code of the software, together with the user-guide manual and list of hardware materials, are publicly available for download at GitHub (https://github.com/CaT-zTools/Deep-CaT-z-Software).

## Acknowledgments

This work was partially financed by FEDER—Fundo Europeu de Desenvolvimento Regional funds through the COMPETE 2020—Operacional Programme for Competitiveness and Internationalisation (POCI), Portugal 2020, and by Portuguese funds through FCT—Fundação para a Ciência e a Tecnologia/Ministério da Ciência, Tecnologia e Ensino Superior in the framework of the projects PTDC/EMD-EMD/31540/2017 (POCI-01-0145- FEDER-031540). We acknowledge the support of the i3S Animal House facility. Ana Gerós was funded by FCT – Fundação para a Ciência e a Tecnologia, grant contract SFRH/BD/137385/2018.

## Author contributions

Ana Gerós: Methodology, Software, Validation, Formal analysis, Visualization, Writing – original draft preparation, Writing – review & editing; Ricardo Cruz: Software, Validation, Formal analysis, Visualization, Writing – review & editing; Fabrice de Chaumont: Writing – review & editing; Jaime S. Cardoso: Methodology, Writing – review & editing; Paulo Aguiar: Conceptualization, Methodology, Writing – review & editing, Supervision.

## Ethics

Animal experimentation: Animal housing and experimental procedures performed according to Portuguese Legislation Dec. Lei n°113/2013 and the European Directive 2010/63/EU on the protection of animals used for scientific purposes. The study was approved by ‘Direção Geral de Alimentação e Veterinária’ (Lisbon, Portugal).

## Supplementary Information

### Supplementary files

**Supplementary Figure 1.**
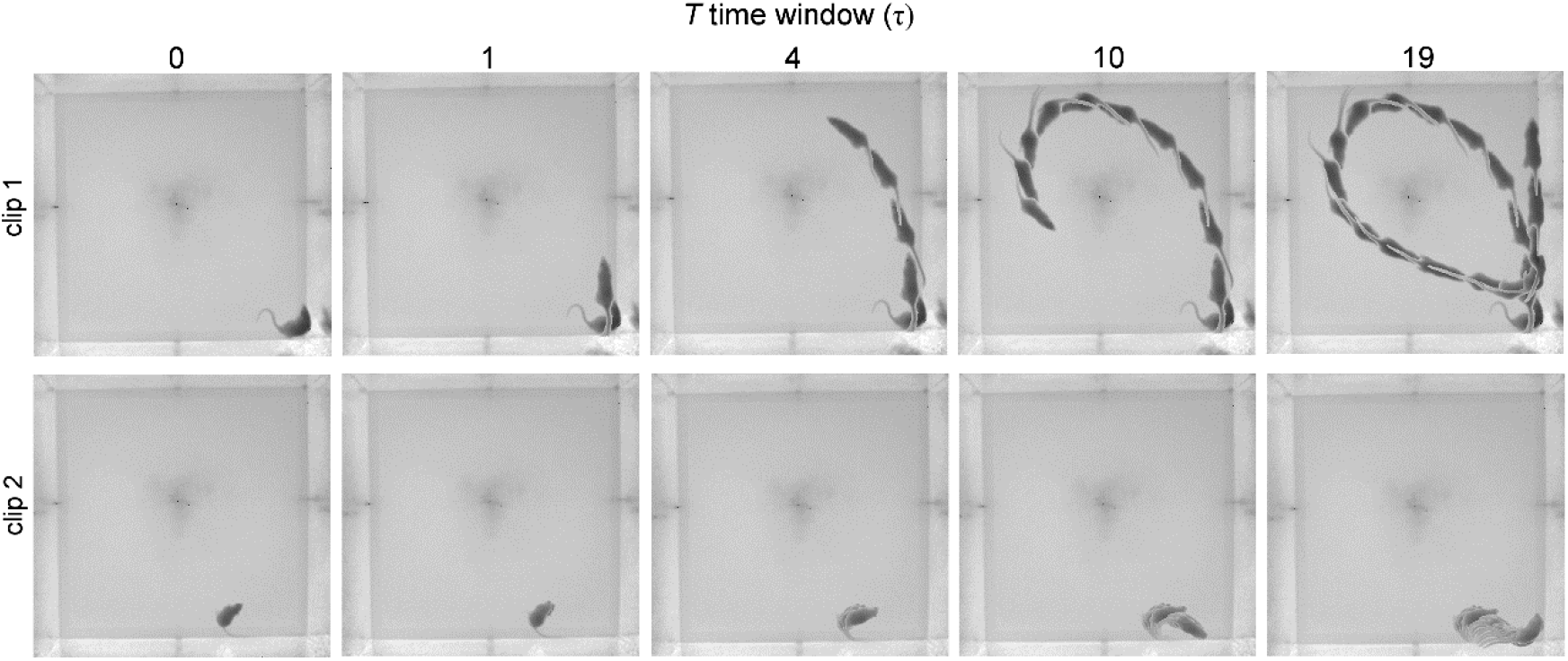
How much temporal information does the network need for rodents’ behavioral learning? Stroboscopic montages in which each animal position represents raw depth frames extracted at every 133 ms, for 2 different *walking* clips and different time windows T, in units of τ (τ = 133 ms). Each stroboscopic image illustrates the depth video sequence input fed to the deep learning network for different values of *T*.

**Supplementary Figure 2.**
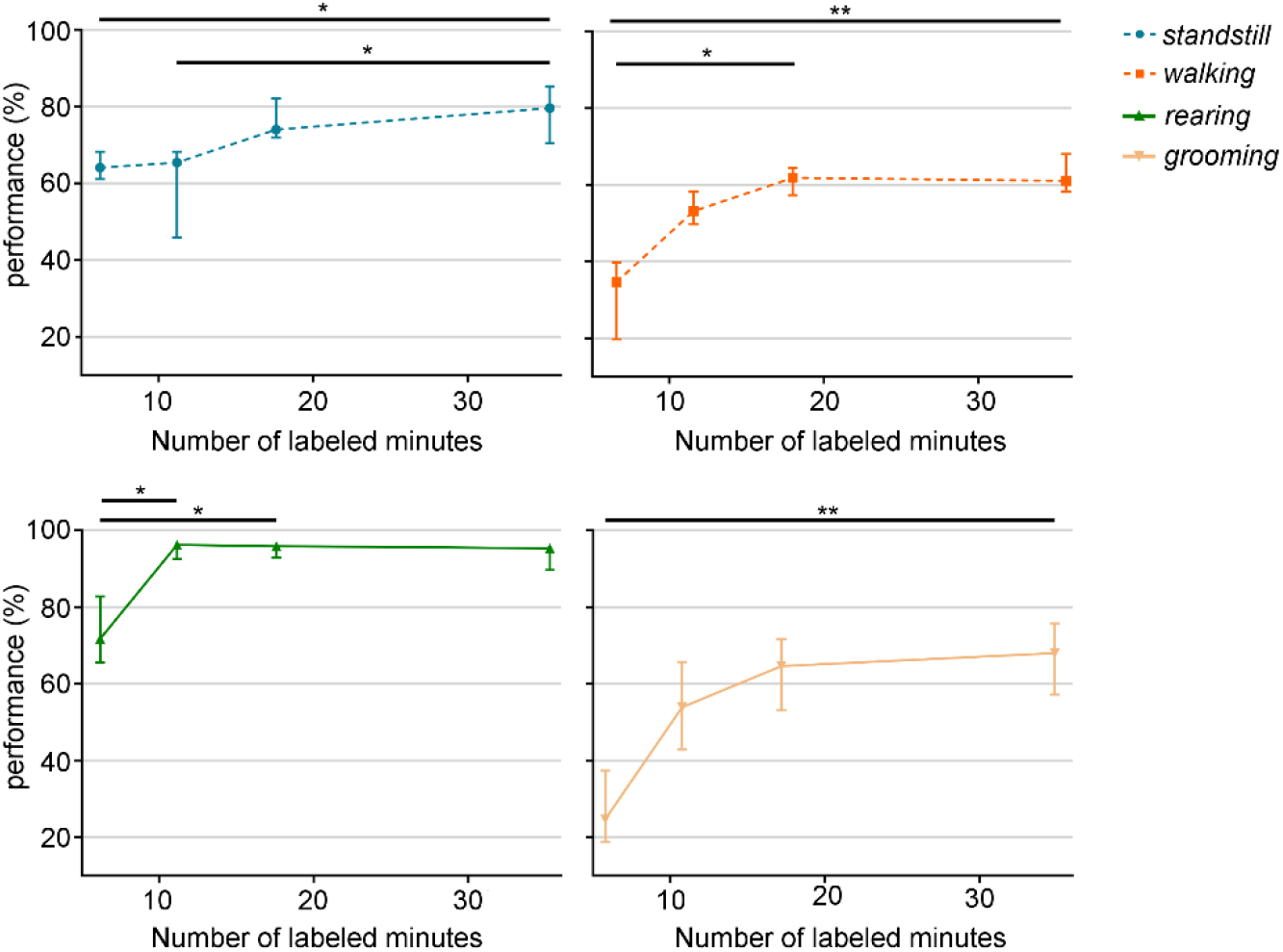
How much information does the network need to learn? Extended statistical analysis for per-class classification performance as function of number of labeled minutes. Data represented as median ± 95% confidence interval (N = 5 trials). * *p* < 0.05; ** *p* < 0.01.

**Supplementary Figure 3.**
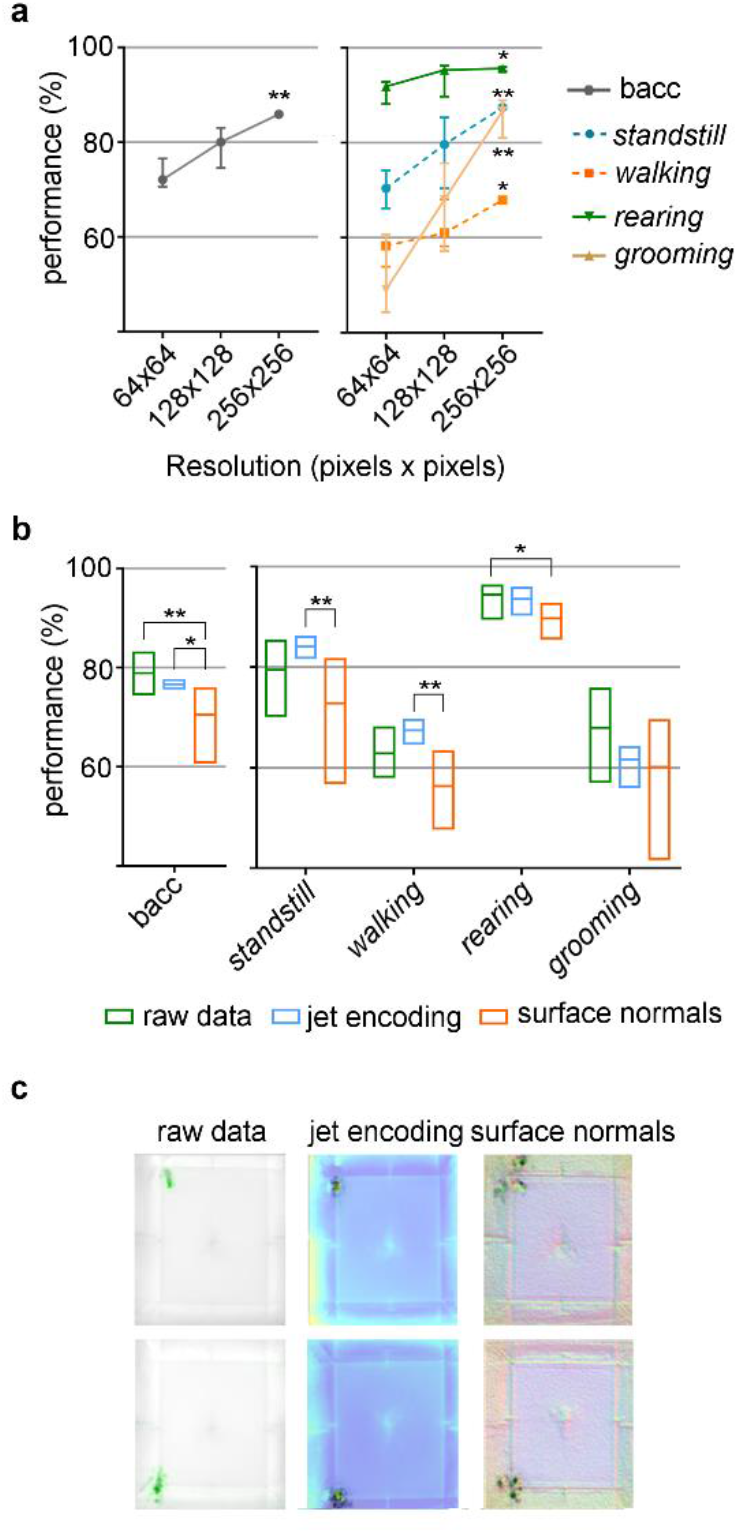
Which input sequence representation is most informative for network’s learning? a. Recognition performance of the single-branch architecture with different input resolutions. * and ** denote statistical significance when compared to the lowest resolution (64x64). **b.** Different depth encodings and corresponding performance, when compared to raw depth input frames. Data represented as median ± 95% confidence interval (N = 5 trials). * *p* < 0.05; ** *p* < 0.01. **c.** Sensitivity analysis for different depth encoding methods (two different frames are shown), with gradients in green or black.

**Supplementary Figure 4.**
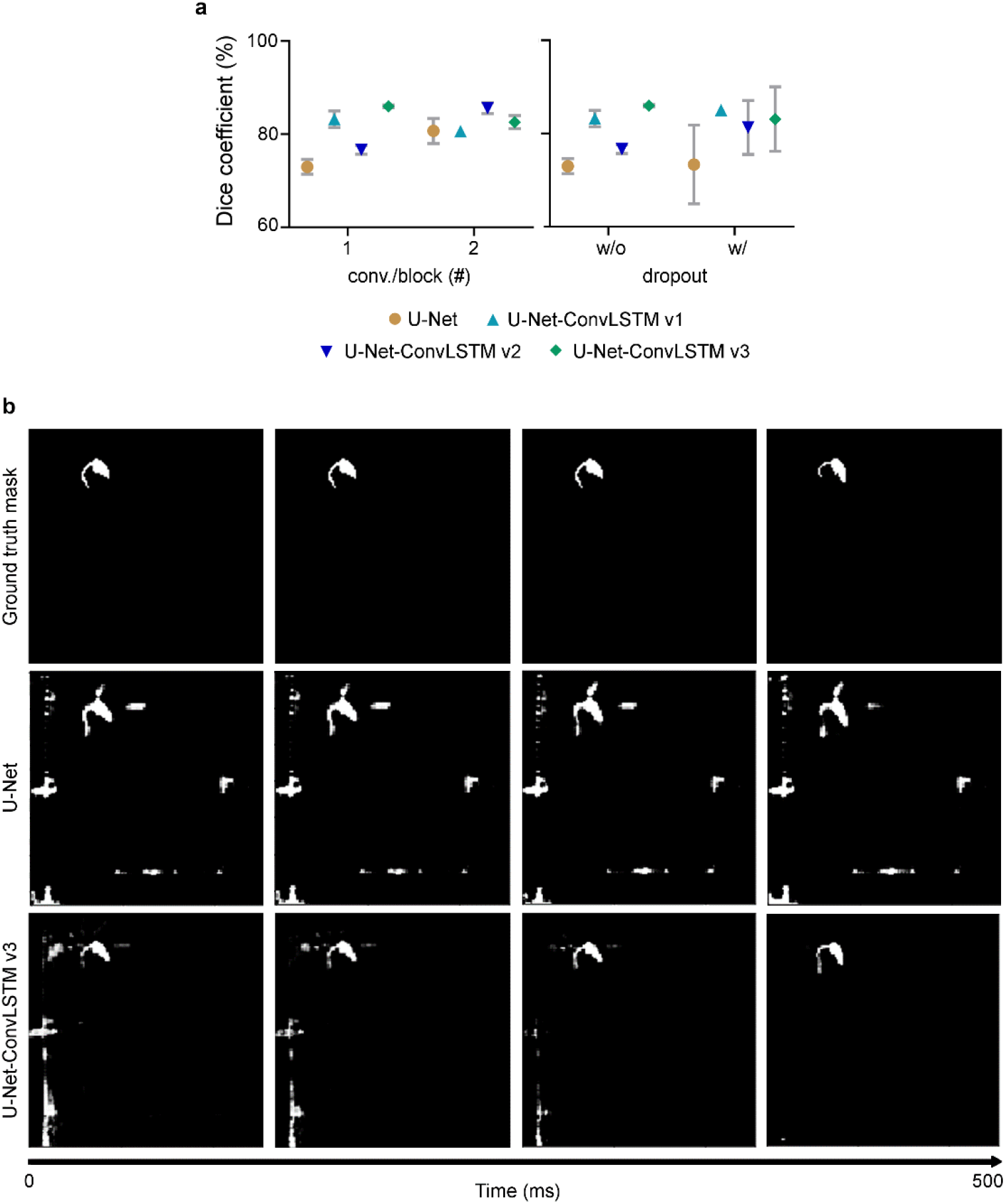
Semantic segmentation results of U-Net-based networks. a. Networks’ performance regarding Dice coefficient for different architectural parameters. Left: number of convolutional layers per block; Right: networks without (w/o) and with (w/) dropout layer at the end of the encoder. The traditional U-Net architecture was extended by placing a convolutional Long Short-Term Memory (ConvLSTM) layer at different positions in the network (U-Net-ConvLSTM), in order to find which position is most suitable for the depth images segmentation task (following Pfeuffer, et al. ^1^ methodology). U-Net-ConvLSTM version 1 (v1) – ConvLSTM layer placed between the encoder and the decoder. U-Net-ConvLSTM version 2 (v2) – ConvLSTM layer placed in the end of the network. U-Net-ConvLSTM version 3 (v3) – a combination of the last two versions. Data represented as median ± 95% confidence interval (N = 2 trials). **b.** Sample clips representing original (top) and predicted segmentation masks by the U-Net (middle) and U-Net-ConvLSTM v3 (bottom) networks, for a time window of 500 ms. Black pixels represent the background predictions and white pixels represent foreground (animal) predictions. During the inference, the presence of ConvLSTM layers improves the segmentation masks over time.

## Additional Results

### Input resolution improves behavioral classification performance

As part of the networks’ study, the effect of input resolution was also examined, keeping the single-branch architecture with default parameters (**Supplementary Figure 3a**). As expected, the highest resolution (256x256) achieved the best results, with an overall performance of 85.9% [82.8 – 86.6]%. All behavioral events seem to benefit from increased resolution, in particular *grooming*, with an increase of approximately 44% over the lowest resolution. The fact that *grooming* events seem to need both higher temporal and spatial resolutions makes it the most sensitive and complex behavior to recognize.

### Raw depth video inputs are the most informative for the learning process

Depth data encodes distance from the sensor to the captured scene and the information of each pixel is of a different nature, when compared to RGB images which were originally directly used as input for the CNNs. Thereby, the questions that arise are will CNNs learn as effectively when using raw depth images without any encoding, and, if not, how should a depth image be encoded to be used as inputs in CNNs so that it can learn more meaningful features for rodents’ classification challenge. Networks were then trained with varying input depth encoding (**Supplementary Figure 3b**). Regarding per-class recognition, the negative effect on network’s learning when using surface normal encoding is even more pronounced. One possible explanation is that when using a colorization method based on the calculation of surface normal, the reflexes on the walls of the open-field during, for example, grooming events (which are always near open-field’s periphery) are more visible and may be interfering with networks’ learning. Sensitivity analysis can be used to identify the most relevant input features during the learning process, by calculating heatmaps from pixel-wise normalized gradients (derivative of class model’s predictions with respect to pixel values). This impact on model’s prediction is exemplified on **Supplementary Figure 3c,** where, by using surface normals, periphery pixels seem to have a stronger influence on model’s prediction (gradient colored as black pixels), when compared to pixels from networks trained with raw depth frames (gradient colored as green pixels). Overall, behavioral learning does not seem to benefit from any of these typical depth input representations.

